# Standardised imaging pipeline for phenotyping mouse laterality defects and associated heart malformations, at multiple scales and multiple stages

**DOI:** 10.1101/516039

**Authors:** Audrey Desgrange, Johanna Lokmer, Carmen Marchiol, Lucile Houyel, Sigolène M. Meilhac

## Abstract

Laterality defects are developmental disorders resulting from aberrant left/right patterning. In the most severe cases, such as in heterotaxy, they are associated with complex malformations of the heart. Advances in understanding the underlying physiopathological mechanisms have been hindered by the lack of a standardised and exhaustive procedure in mouse models, for phenotyping left/right asymmetries of all visceral organs. Here, we have developed a multimodality imaging pipeline, which combines non-invasive micro-ultrasound imaging, micro-CT and HREM, to acquire 3D images at multiple stages of development and at multiple scales. Based on the position in the uterine horns, we track, in a single individual, the progression of organ asymmetry, the *situs* of all visceral organs in their thoracic or abdominal environment, together with fine anatomical left/right asymmetries of cardiac segments. We provide reference anatomical images and organ reconstructions in the mouse, and discuss differences with humans. This standardised pipeline, which we validated in a mouse model of heterotaxy, offers a fast and easy-to-implement framework. The extensive 3D phenotyping of organ asymmetry in the mouse uses the clinical nomenclature for direct comparison with patient phenotypes. It is compatible with automated and quantitative image analyses, which is essential to compare mutant phenotypes with incomplete penetrance and gain mechanistic insight into laterality defects.

**Summary statement:** Laterality defects, which combine anomalies in several visceral organs, are challenging to phenotype. We have now developed a standardised approach for multimodality 3D imaging in mice, generating quantifiable phenotypes.

## Introduction

Laterality defects are developmental disorders, caused by impaired left-right patterning. They affect collectively up to 1/2,000 live births, and comprise a spectrum of malformations ranging from asymptomatic situs inversus or dextrocardia to severe heterotaxy (Desgrange et al., 2018; Lin et al., 2014). Heterotaxy corresponds to abnormal symmetry of the viscera (isomerism), and/or situs discordance between visceral organs (Van Praagh, 2006). Many visceral organs (lung, spleen, stomach, intestine, liver and pancreas) may be targeted and functionally impaired, with an abnormal position in the abdominal or thoracic cavity (situs) or an impaired asymmetric shape. Diagnosis is made on the combination of 3 out of 8 criteria, including abdominal situs abnormality, spleen abnormality, isomerism of bronchi and the lungs, biliary atresia, intestinal malrotation, congenital heart defects, isomerism of the atrial appendages, systemic venous anomalies (Lin et al., 2014). The prognosis of heterotaxy mainly depends on the cardiac malformation, which can be complex, with profound functional impacts, such as abnormal oxygen supply or obstructed blood flow. Diagnosis of complex congenital heart defects is performed using the segmental approach developed by Van Praagh (Van Praagh, 1972), analysing separately the atria, the ventricles and the great arteries. Anatomical features are used to differentiate each cardiac segment (left/right ventricle (Van Praagh and Van Praagh), left/right atrium (Uemura et al., 1995)), whereas the heart phenotype is based on the position, in the thoracic cavity, of the morphological left and right cardiac segments and their connection relative to each other (Jacobs et al., 2007).

In the last two decades, experiments in the mouse/animal model have provided insight into how left-right patterning is established in the early embryo (Hamada and Tam, 2014), from the activity of a left-right organiser, in which fluid flow generated by motile cilia plays a central role (Nonaka et al., 1998). Heart looping at E8.5 is the first morphological sign of left/right asymmetric morphogenesis (Desgrange et al., 2018). During this process, the heart tube, which is initially straight, with the right ventricle lying cranially to the left ventricle, becomes helical. This establishes the relative position of the different cardiac segments, such that the right ventricle lies on the right side of the left ventricle (Le Garrec et al., 2017). Thus, the definitive left/right position of organ segments is the result of asymmetric morphogenesis. In contrast, gastrulation marks a physical separation, on either side of the primitive streak, between left and right precursor cells, which receive asymmetric signalling, as for example from the left determinant Nodal (Collignon et al., 1996). Left and right precursor cells can be traced to assess how they contribute to different regions of an organ. DiI labelling and clonal analyses have thus provided insight into the left/right embryological origin of liver lobes (Weiss et al., 2016) or cardiac segments (Domínguez et al., 2012; Lescroart et al., 2010; Lescroart et al., 2012). However, the mechanisms of asymmetric organ morphogenesis remain largely unknown.

The mouse provides a good model for the study of laterality defects, given the array of genetics tools available to reproduce genetic alterations, and given anatomical similarities in mammals. However, there are also anatomical variations between the mouse and human, which can be extracted from fragmented analyses of individual mouse organs (Ciszek et al., 2007; Fiebig et al., 2012; Henke et al., 2018; Kaufman and Richardson, 2005; Nguyen et al., 2015; Thiesse et al., 2010; Webb et al., 1996). Thus, a comprehensive description of laterality features in all mouse visceral organs, and its relevance to clinical diagnosis has been lacking. Analyses of mouse mutant lines have been limited by several aspects. Mutations of genes involved in the left-right organiser lead to several categories of phenotypes, which are not fully penetrant and often observed with a randomised frequency. This is the case for mutations impairing ciliogenesis, which are associated with randomised heart looping direction in the embryo and congenital heart defects at birth (Layton, 1976; Vierkotten et al., 2007). Phenotype randomisation requires observations in a high number of individuals and hinders correlation of phenotypes at different stages. The phenotype description is often incomplete, focusing on a few organs of interest and with quantification in a small sample size. Phenotypes can be described in invasive open chest dissections or explanted organs, thus limiting conclusions on organ *situs*. Alternatively, complex malformations are diagnosed in 2D histological sections, which are associated with tissue distortion and may miss important features without the third dimension and the continuity of structures. In addition, phenotypes described by developmental biologists do not always refer to the clinical nomenclature, hence limiting the possibility of cross-correlations with patient phenotypes. Thus, understanding the origin of laterality defects have been hindered by the lack of a standardised and exhaustive procedure of phenotyping. Advances in 3D volumetric imaging have opened the possibility to image the non-transparent mouse organism, at a smaller scale compared to humans. *In utero* mouse development is now accessible by high-frequency micro-ultrasound imaging (Foster et al., 2011) or Optical Coherence Tomography (OCT) (Syed et al., 2011). With other approaches, such as Optical Projection Tomography (OPT) (Sharpe et al., 2002), OCT (Lopez et al., 2015) or X-ray micro-Computed Tomography (micro-CT) (Degenhardt et al., 2010), the structure of internal organs can be reconstructed. With a higher imaging depth and wider field of view, micro-CT was selected for routine screening of mouse mutants (Wong et al., 2014). Finally, histological based approaches such as Episcopic Fluorescence Image Capture (EFIC) or High Resolution Episcopic Microscopy (HREM) (Mohun and Weninger, 2012) reach very high spatial resolution, able to resolve subtle anatomical variations, as exemplified in the heart (Geyer et al., 2017). A challenge to fully describe laterality defects, is to combine images at multiple scales, to resolve the *situs* of all visceral organs together with fine anatomical left/right features, and at multiple stages of development, to understand the origin of the defects.

Here, we report a novel multimodality imaging pipeline to phenotype laterality defects in 3D in the mouse. To assess the shape of the embryonic heart loop *in vivo*, we first perform non-invasive micro-ultrasound imaging on a pregnant mouse. By recording the position of each embryo in the uterine horns, correlations with another stage of development is possible. Each individual is thus tracked just before birth, at E18.5 by micro-CT, to determine the *situs* of visceral organs in their endogenous environment, i.e. without dissection, as well as to resolve vascular connections. Finally, to evaluate the fine cardiac anatomy, including left/right features, we acquire images of the isolated E18.5 heart by HREM. Based on sequential established imaging modalities, the novel imaging pipeline is a simple and standardised procedure for an extensive phenotyping of organ laterality in the mouse. To guide phenotyping, we provide state-of-the-art reference images annotated with the clinical nomenclature.

## Results

### Imaging the embryonic heart loop *in utero* by micro-ultrasound imaging

Heart looping is the first morphological sign of left/right asymmetry in the developing embryo and anomalies in this process are associated with congenital heart defects (see Desgrange et al., 2018). To be able to correlate the shape of the heart at two different stages, imaging within the same individual is required. As a first step of an imaging pipeline (Fig. 1), we used non-invasive micro-ultrasound imaging to evaluate the shape of the embryonic heart loop, without perturbing embryo development. We selected E9.5 as a stage when heart looping is complete (Fig. 2A), and used 3D reconstruction of heart shape in fixed samples (Fig. 2B, Movie 1) as a framework of analysis of lower-resolution images. We show that we can identify, in a pregnant mouse, individual embryos, each within a deciduum, by micro-ultrasound imaging. Each embryo is numbered according to its position in the uterine horns (Fig. 1A). At necropsy 9 days after imaging, the number of E18.5 fetuses found in each uterine horn was consistent (6 and 3 in litter #1, 3 and 5 in litter #2, 4 and 3 in litter #3, 4 and 4 in litter #4, respectively on the left and right). 31/32 fetuses were collected alive, in accordance with the good viability of embryos after micro-ultrasound imaging. For a standardised analysis of the embryonic heart shape, a projection of the 3D micro-ultrasound images was generated, and coronal tissue sections extracted, independently of the orientation of image acquisition. In coronal sections, we used the head and tail as landmarks of the cranio-caudal axis of the embryo, and the heart and neural tube for its ventral-dorsal axis (Movie 2). Based on these two axes, the orientation of the left/right axis of the embryo can be determined (Fig. 2 C-D). At E9.5, the fast heartbeats facilitate the localisation of the heart tube (Movie 2). The analysis of the shape of the embryonic heart, is based on its organisation in distinct regions, positioned sequentially along the axis of the cardiac tube (Le Garrec et al., 2017). The outflow tract can be identified as the connection of the heart tube to the cranio-dorsal part of the embryo, whereas the right and left ventricles follow ventrally, separated by a sulcus, on the right and left of wild-type embryos respectively (Fig. 2C), compared to the atria, which are more dorsal and caudal, also as two chambers on the right and left of wild-type embryos (Fig. 2D). Thus, we show that micro-ultrasound imaging is an appropriate and non-invasive method to assess the overall shape of the embryonic heart *in vivo*, as early as E9.5.

**Figure 1:**
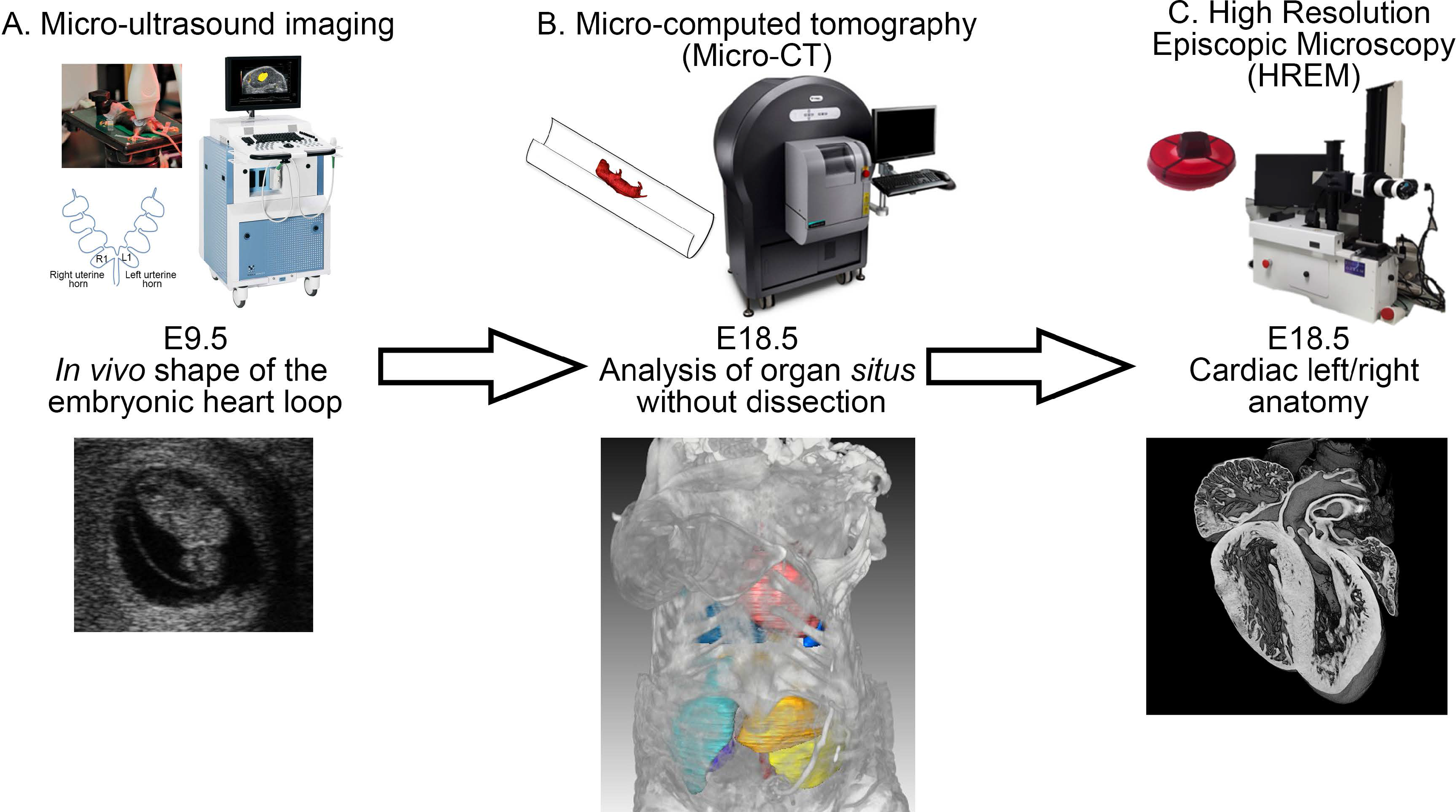
Multimodality imaging pipeline of left/right asymmetries in the mouse. (A) Micro-ultrasound imaging of a pregnant mouse to assess *in vivo* the shape of the embryonic heart loop at E9.5. The position of the embryo in the uterine horn is recorded as schematised, L1 and R1 being the first embryo next to the vagina, in the left and right horns respectively. (B) Micro-computed tomography (micro-CT) of the same individuals at E18.5, to image the *situs* of thoracic and abdominal organs. (C) High Resolution Episcopic Microscopy (HREM) on the explanted E18.5 heart to image its left/right anatomical features. In each panel, the sample preparation is shown on the top left, the equipment on the top right, and an example of an image on the bottom (A, optical section; B, 3D projection with segmented organ contours ; C, 3D projection).

**Figure 2:**
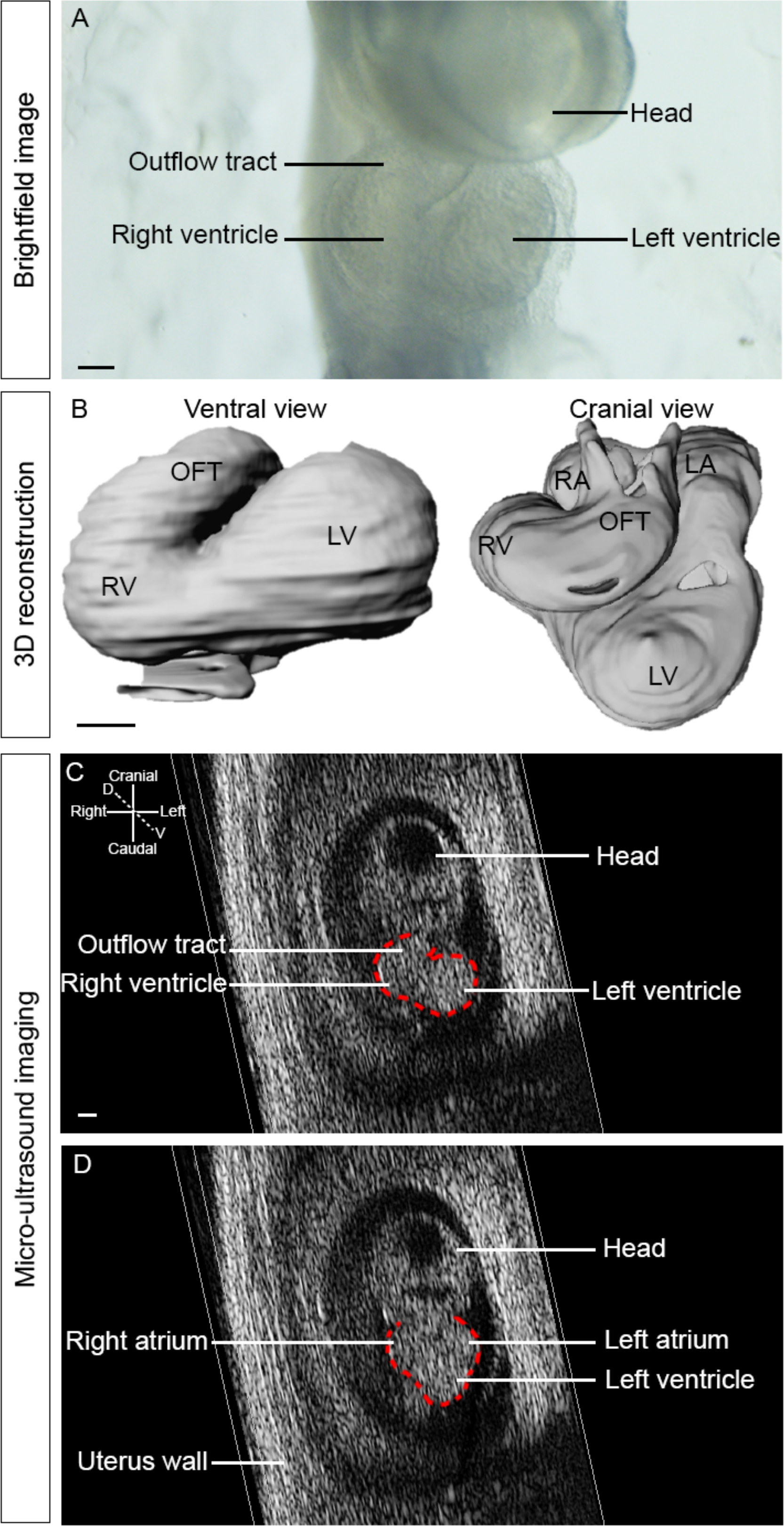
Micro-ultrasound imaging of the embryonic heart loop *in utero*. (A) brightfield image of an explanted embryo at E9.5, showing the shape of the embryonic heart. (B) 3D reconstruction of the embryonic heart shape on a ventral and cranial view, after segmentation of HREM images at E9.5 showing the helical shape of the heart. Ventral (C) and dorsal (D) sections of a 3D+t image of a E9.5 wild-type embryo acquired by micro-ultrasound imaging *in utero*. The contour of the embryonic heart is outlined in red. D, dorsal; LV, Left ventricle; OFT, Outflow tract; RV, Right ventricle, V, ventral. Scale bar: 100 μm

### Imaging the *situs* of thoracic and abdominal organs in fetuses by micro-CT

To assess the *situs* of all visceral organs, which can potentially be abnormal in laterality defects, a rapid imaging procedure, resolving several organs inside the body was required. Micro-computed tomography, with a field of 10×10×10 mm, can acquire in 3 min images of all thoracic and abdominal organs, without any dissection of the fetus or neonate mouse. To be able to correlate phenotypes at birth with images of the embryo, samples were collected at E18.5, just before birth, when their position in the uterus can be tracked. Micro-CT provides 3D images, which can be segmented to reconstruct organ shapes in 3D, or can be re-sectioned optically in any relevant orientation.

Micro-CT scans of E18.5 fetuses were analysed for the distinct left/right features of visceral organs, including the lung, liver, stomach, colon and spleen (Fig. 3). As previously described (Thiesse et al., 2010), the right bronchus was detected from its first division, which is more cranial compared to that of the left bronchus (Fig. 3A). In the mouse, the right lung is divided into four lobes (Thiesse et al., 2010), which are clearly identified in micro-CT scans : the right superior lung lobe (RSLuL) which is more cranial, the right middle lung lobe (RMLuL) which is more dorsal and caudal to the RSLuL, the right inferior lung lobe (RILuL) which is more caudal and ventral, and the post caval lung lobe (PCLuL) which is smaller and located more medially, dorsal to the heart. In contrast, the left lung is composed of a single, large lobe, the left lung lobe (LLuL) (Fig. 3A, E). The mouse liver is also a bilateral and asymmetric organ, located in the abdominal cavity abutting the diaphragm. All liver lobes (Fiebig et al., 2012) can be identified in micro-CT scans (Fig. 3B-C, E). There are two lobes on the left, with the left medial liver lobe (LMLiL) that is more cranial and smaller than the left lateral liver lobe (LLLiL) lying over the stomach. The right liver is subdivided into four lobes. The right medial liver lobe (RMLiL) is cranial and on the right of the gall bladder, and the right lateral medial liver lobe (RLLiL) is under the RMLiL and smaller. The right caudal liver lobe (RCLiL) is more caudal abutting the right kidney. Finally, the papillary process (PP) is located between the stomach and the RLLiL, wrapping around the oesophagus. Despite its medial location, clonal analyses indicate that it is embryologically more closely related to the right lobes and thus can be considered as a right structure (Weiss et al., 2016). Other visceral organs, with an asymmetric position and shape were detected in micro-CT scans (Fig. 3D-E). As in humans, the stomach is located on the left under the LLLiL and the spleen runs along it caudally. The colon, as reported previously (Henke et al., 2018; Nguyen et al., 2015), has a short C-shape curvature, going dorsally and on the left. Thus, micro-CT imaging is a simple and powerful technique to resolve the laterality features of visceral organs, both qualitatively (organ position, asymmetric shape) and quantitatively (organ size, number of lobes). Here, the left and right nomenclature is mainly based on the position within the thoracic or abdominal cavity, with the exception of the PCLuL, connected with the right bronchus, and the papillary process, clonally related to the right liver lobes. We provide annotated 3D reconstructions of organ shape in their endogenous configuration within the body (Movie 3), which will be useful to phenotype mouse models of laterality defects.

**Figure 3:**
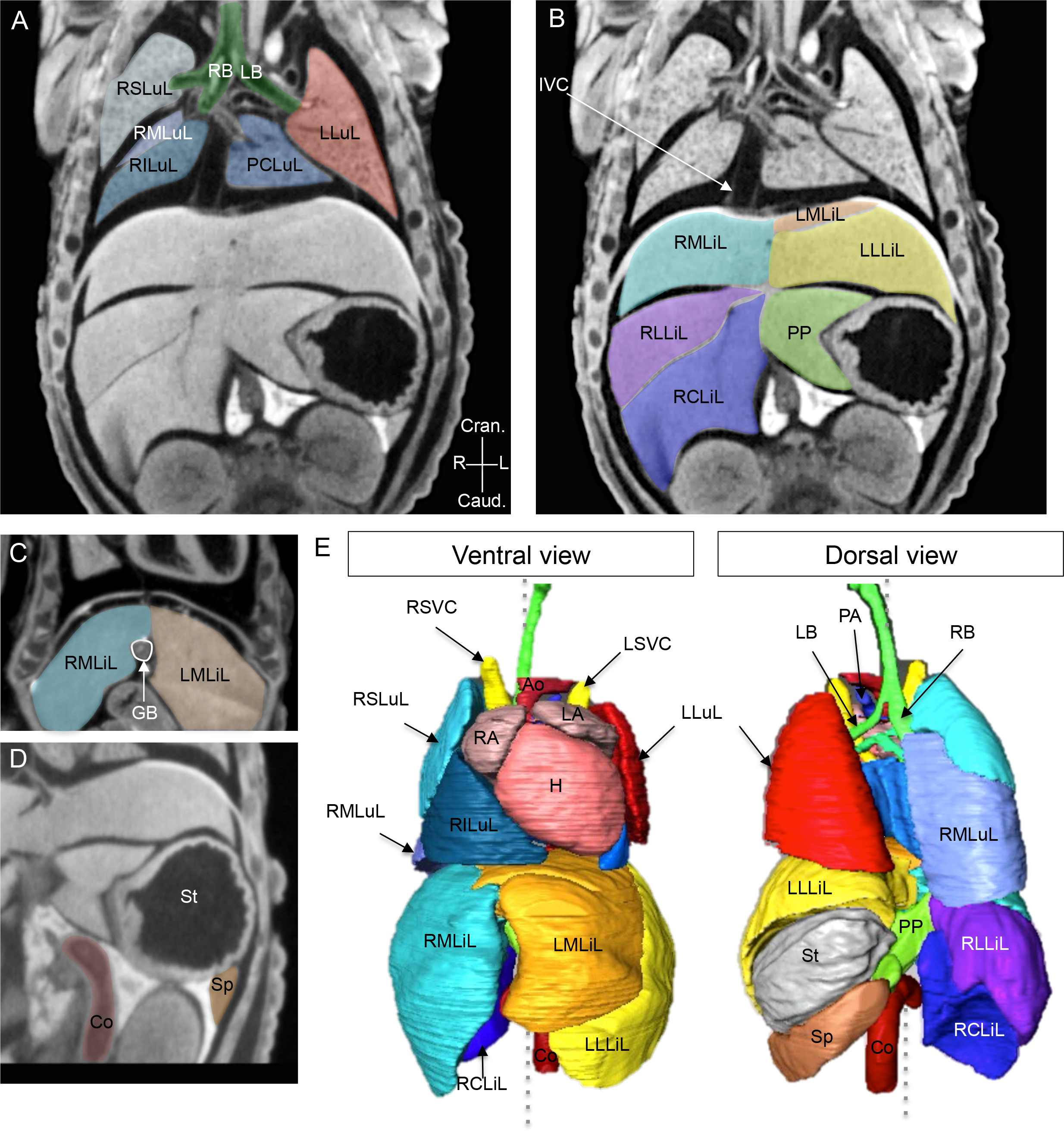
Imaging of the *situs* of thoracic and abdominal organs by micro-CT at E18.5. (A-D) Coronal sections from 3D images, acquired by micro-CT, of wild-type fetuses at E18.5, showing the thoracic and abdominal organs, highlighted in different colours. (E) 3D reconstruction of the shape and position of organs within the body, from a ventral or dorsal view. The grey dotted line represents the plane of bilateral symmetry. Ao, aorta; Co, colon; Cran., cranial; Caud., caudal; GB, gall bladder; H, heart; IVC, inferior *vena cava*; L, left; LA, left atrium; LB, left bronchus; LLLiL, left lateral liver lobe; LLuL, left lung lobe; LMLiL, left medial liver lobe; LSVC, left superior *vena cava*; PA, pulmonary artery; PCLuL, post-caval (right) lung lobe; PP, papillary process; R, right; RA, right atrium; RB, right bronchus; RCLiL, right caudal liver lobe; RILuL, right inferior lung lobe; RLLiL, right lateral liver lobe; RMLiL, right medial liver lobe; RMLuL, right middle lung lobe; RSLuL, right superior lung lobe; RSVC, right superior *vena cava*; Sp, spleen; St, stomach.

### Imaging the *situs* of the heart and its connections with the great vessels by micro-CT

As for other visceral organs, the *situs* of the heart is detectable in micro-CT scans, based on the location of the apex, which normally points toward the left, a situation referred to as levocardia (Fig. 4A-B). Micro-CT also provides sufficient resolution to analyse the great vessels, which are asymmetric structures. In healthy patients, cardiovascular structures are named or referred to as left or right (shown in red and blue respectively in Fig. 4), on the basis of their contribution to the systemic and pulmonary blood circulations, which are driven by the left and right chambers, respectively. The aorta forms an arch toward the left side of the trachea and oesophagus (Fig. 4A-D), and descends on the left of the thoracic cavity (Fig. 4E-F). It is connected to the left ventricle (Fig. 4B), whereas the pulmonary artery, which branches closer to the heart, is connected to the right ventricle. The four pulmonary veins, from the left and right lung lobes, are fused into a single collector that connects to the left atrium (Fig. 4F). The inferior *vena cava* runs on the right of the body, from abdominal organs to the vestibule of the right atrium (Fig. 4A, C). In the mouse, the right and left superior *vena cava*, both containing blood from the systemic circulation, arrive from the right and left of the head respectively, and connect to different regions of the right atrium, the roof and vestibule respectively (Fig. 4A-B, D). Thus, by providing a scan of the entire thoracic and abdominal cavities, micro-CT is able to resolve not only the *situs* of visceral organs, but also the complex relative position and connections of the great vessels, which are important features in laterality defects.

**Figure 4:**
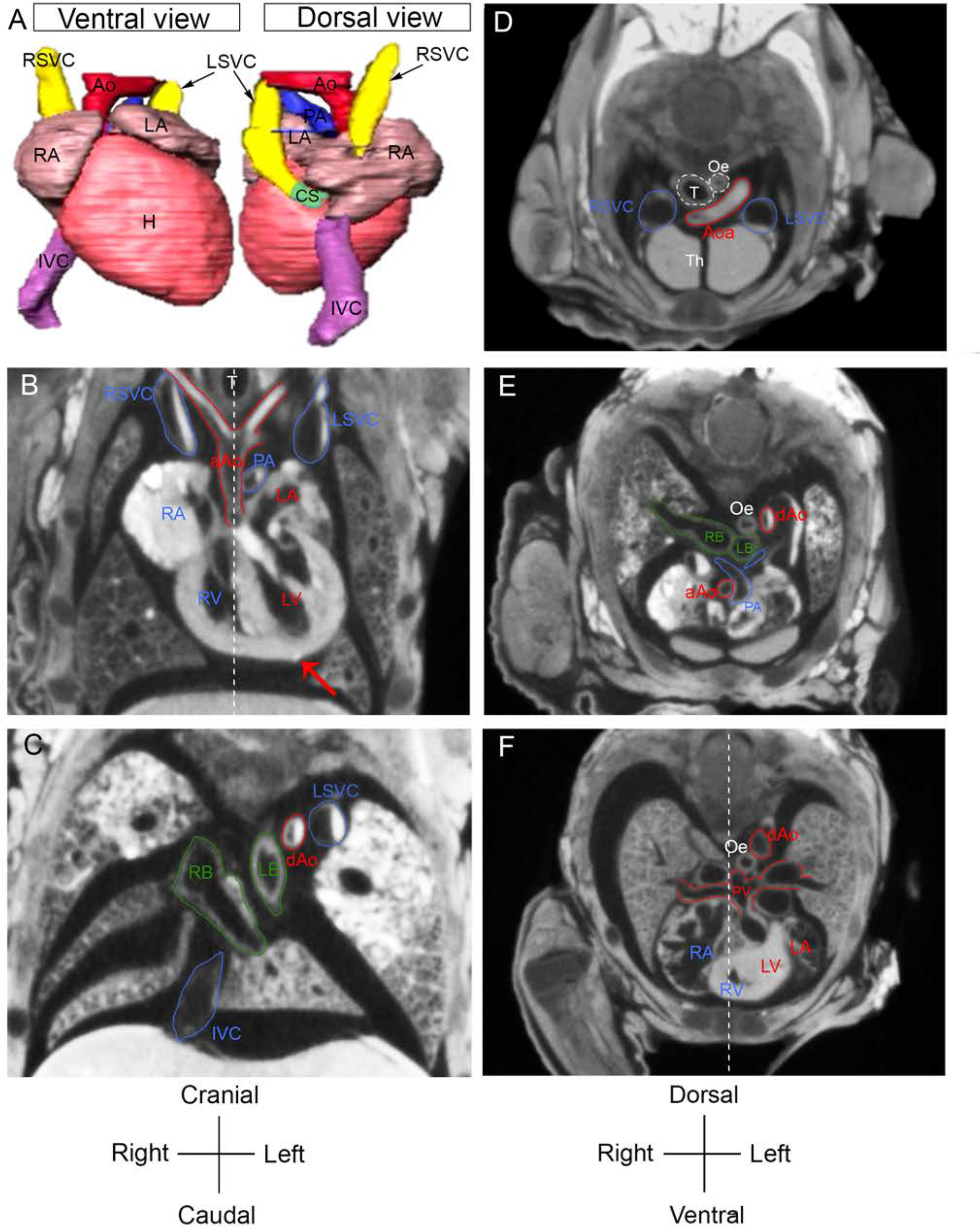
Imaging of the *situs* of the heart and the great vessels by micro-CT. (A) Ventral and dorsal views of a 3D reconstruction of the heart (H) and great vessels, after image segmentation. Coronal (B-C) and transverse (D-F), in a cranio-caudal order) optical sections from micro-CT scans of E18.5 fetuses, showing the *situs* of the heart and the great vessels. The red arrow points to the heart apex, which is on the left (levocardia) in a wild-type sample. The white dotted line represents the plane of bilateral symmetry, bisecting the neural tube. Structures specific to the circulation of deoxygenated and oxygenated blood are annotated in blue and red, respectively ; bronchi are in green, other structures in white. aAo, ascending aorta; Aoa, aortic arch; CS, coronary sinus; DA, ductus arteriosus; dAo, descending aorta; IVC, inferior *vena cava*; LA, left atrium; LB, left bronchus; LV, left ventricle; LSVC, left superior *vena cava*; Oe, oesophagus; PA, pulmonary artery; PV, pulmonary veins; RA, right atrium; RB, right bronchus; RSVC, right superior *vena cava*; RV, right ventricle; T, trachea; Th, thymus.

### Imaging fine anatomical asymmetries in the heart by HREM

In the clinics, phenotyping the left-right asymmetry of the heart is based on a segmental approach (Van Praagh, 1972), analysing separately the fine anatomical features of the two atria, the two ventricles and the two great arteries, beyond their position within the thoracic cavity. This requires a histological resolution higher than that of micro-CT, which is reached by HREM of the explanted E18.5 heart, providing 3D images, with a resolution below 4 μm. Such 3D images can be exploited to generate 3D projections of the heart, as well as histological sections in any orientation relevant to identify distinct left-right features.

The left and right identity of the atria can be distinguished based on their appendages (Uemura et al., 1995). The right atrial appendage has pectinate muscles which extend on the entire atrial chamber, including at the exit of the inferior *vena cava* (asterisk Fig. 5A-B). In contrast, the left atrial appendage corresponds to more confined pectinate muscles at the tip of the atrial chamber (Fig. 5A-B), whereas the vestibular region is smooth. Another asymmetric feature is the specific connection of the right atrium with the inferior caval vein (Van Praagh, 1992) at the level of the Eustachian valve (Fig. 5A-B). In the mouse, the coronary sinus, which lies in continuity with the left superior caval vein (Fig. 4A), receives blood from the coronary veins and opens into the right atrium (Fig. 5C-D). The correct configuration, with a morphologically left atrium on the left of the thoracic cavity and the morphologically right atrium on the right, is described as *situs solitus* (S).

**Figure 5:**
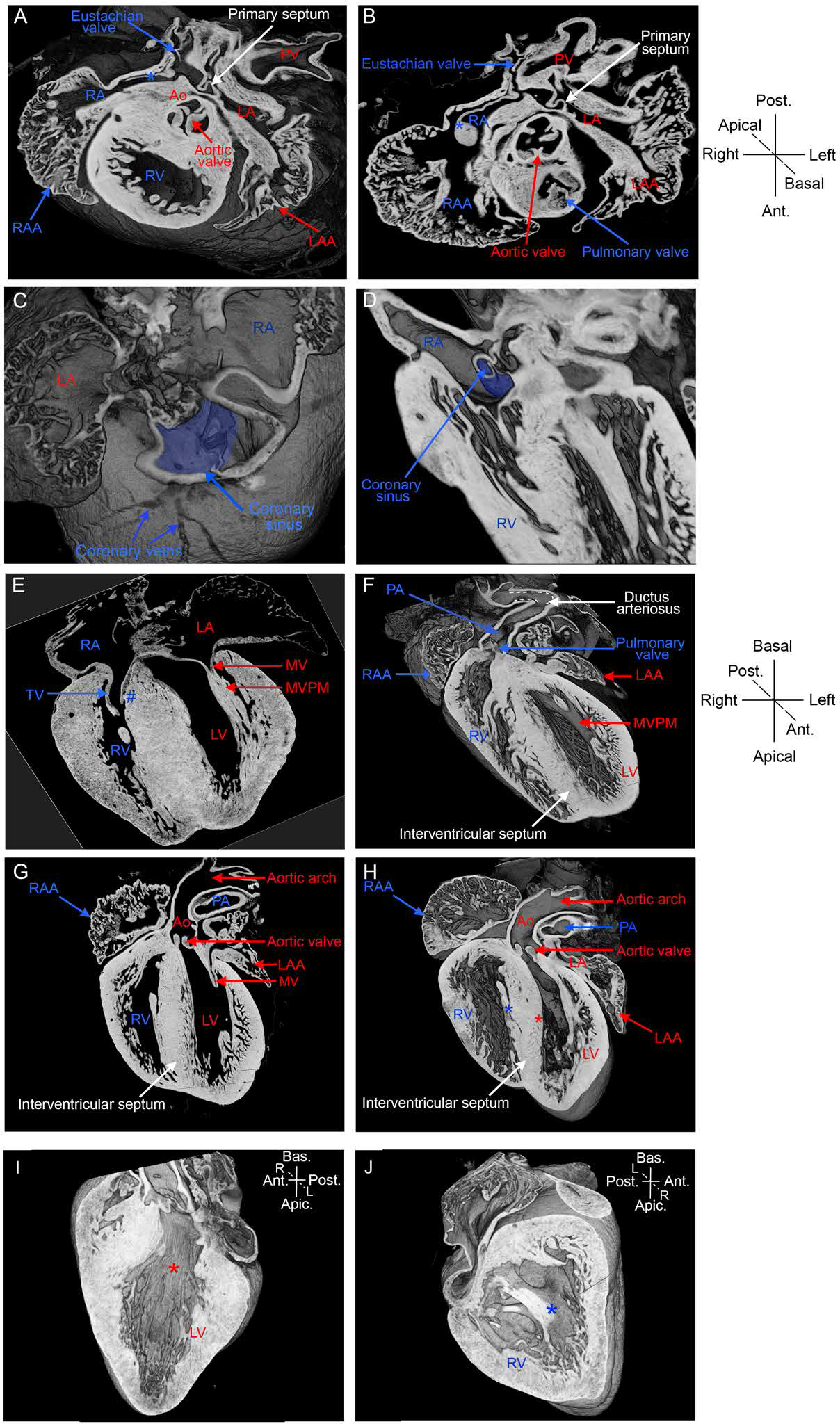
Imaging the fine cardiac anatomy by HREM. 3D projection (A) and transverse section (**B**) of E18.5 hearts, imaged by HREM, at the level of the semilunar valves, showing the relative positions of the aorta and pulmonary artery, as well as the differential distribution of pectinate muscles in the left and right atrial chambers. The blue asterisk points to a pectinate muscle of the right atrial appendage, which attaches to the Eustachian valve, at the end of the inferior *vena cava*. (C-D) 3D projections showing the coronary sinus, highlighted in blue, collecting the coronary veins and connected to the right atrium. 3D projections (E, G) and coronal sections (F, H) showing the cardiac chambers, the great arteries and the valves. (I-J) 3D projections of the interventricular septum showing the smooth basal surface in the left ventricle (red asterisk, I), and trabeculated surface in the right ventricle (blue asterisk, J). The tricuspid valve has one septal leaflet (blue #). Structures specific to the circulations of deoxygenated or oxygenated blood are annotated in blue and red, respectively; other structures are in white. Apic., apical; Ant., anterior; Ao, aorta; Bas., basal; L, left; LA, left atrium; LAA, left atrial appendage; LV, left ventricle; MV, mitral valve; MVPM, mitral valve papillary muscles; PA, pulmonary artery; Post., posterior; R, right; RA, right atrium; RAA, right atrial appendage; RV, right ventricle; TV, tricuspid valve.

The right and left ventricles can be identified based on their atrio-ventricular valves and the trabeculation of the interventricular septum, at a basal level. At the entrance of the right ventricle, the tricuspid valve is more apical compared to the mitral valve of the left ventricle (Fig. 5E). The tricuspid valve has a septal attachment, in addition to papillary muscles, whereas the mitral valve has papillary muscles and no septal attachment (Fig. 5E-G). The left ventricle has a smooth septal surface and fine apical trabeculations (Fig. 5H-I), whereas the right ventricle has a trabeculated septal surface with coarse apical trabeculations (Fig. 5H, J). The correct configuration, with the morphologically left ventricle on the left of the thoracic cavity and the morphologically right ventricle on the right, is referred to as a D-Loop (D).

In keeping with the rightward rotation of the outflow tract during development (Bajolle et al., 2006; Le Garrec et al., 2017), the position of the great arteries is also a manifestation of left-right asymmetry. The aorta, defined from its connection to the head, normally arises from the left ventricle (Fig. 5G-H) and crosses the pulmonary artery, defined from its connection to the lung, and normally arising from the right ventricle (Fig. 5F). The aortic arch is oriented toward the left of the body (Fig. 5G-H). In transverse sections of the heart, at the level of the aortic and pulmonary valves, the aorta is positioned dorsally and on the right compared to the pulmonary artery (Fig. 5B). In the fetus, the aorta is connected with the pulmonary artery via the ductus arteriosus (Fig. 5F), which will regress after birth. This correct configuration of the great arteries, in the segmental analysis of the heart structure, is referred to as *situs solitus* (S).

With the segmental approach, the nomenclature of a well-formed heart is thus {S,D,S}, in reference to the position of the atria (*situs solitus*), the ventricles (D-Loop) and the great arteries (*situs solitus*) respectively. This clinical nomenclature can be applied to the mouse. Another descriptor of the correct alignment of cardiac chambers is atrio-ventricular concordance, when the morphological right atrium connects with the right ventricle, and vice versa for the left.

In conclusion, HREM provides a high-resolution 3D image of the mouse heart, which is relevant to the phenotyping of the anatomical left/right differences of cardiac chambers. Together with ultrasound imaging and micro-CT, it provides an extensive phenotyping of the *situs* and asymmetry of visceral organs, including the cardiovascular system.

### Application of the pipeline to phenotype heterotaxy in Rpgrip1l *mutants*

We applied our novel imaging pipeline to a previously characterized mouse model of the heterotaxy syndrome (Delous et al., 2007; Vierkotten et al., 2007). *Rpgrip1l* encodes a ciliary protein localized to the basal body of cilia. It is required for ciliogenesis, including in the left-right organiser, the node. We performed micro-ultrasound imaging of 2 independent litters, corresponding to a total of 18 embryos at E9.5 (Fig. S1). In 9 embryos, we observed the characteristic shape of the embryonic heart loop (Fig. 6A1). In contrast, 5 embryos showed aberrant embryonic heart shapes, including reversed looping (Fig. 6B1) or a straight heart tube with pericardial effusion (Fig. 6C1), and 4 embryos were underdeveloped or with no heartbeat. This is consistent with previous observations, showing anomalies of heart looping in *Rpgrip1l* mutants, in association with bilateral expression of *Pitx2*, a target of the left determinant Nodal (Vierkotten et al., 2007). As expected from the lethality of *Rpgrip1l^−/−^* mutants *in utero* (Vierkotten et al., 2007), 5/18 fetuses at E18.5 were not recovered and 1/18 was underdeveloped and dead. The degenerated decidua and dead fetus were found in the position of embryos with embryonic anomalies. Genotyping by PCR in surviving individuals confirmed the homozygous mutation in fetuses which had abnormal embryonic heart shapes and the absence of it in the normal samples, except for one of the underdeveloped embryos which was heterozygous (Fig. S1).

**Figure 6:**
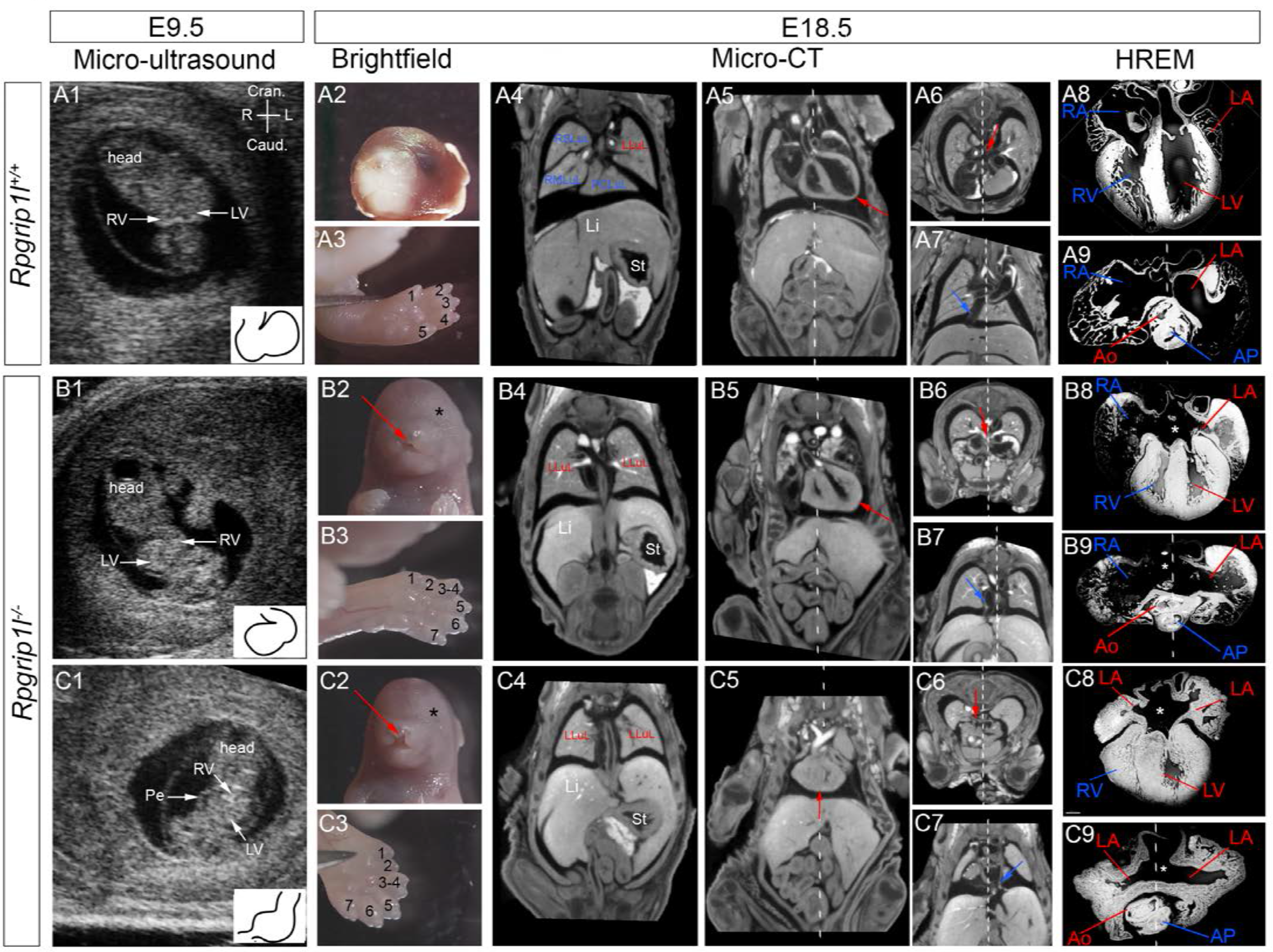
Application of the pipeline to phenotype heterotaxy : example of *Rpgrip1l* mutants. (A1-C1) Representative section of the 3D+t dataset from micro-ultrasound imaging of *Rpgrip1l*^*+/+*^(A1) and *Rpgrip1l*^*−/−*^ (B1-C1) embryos at E9.5. The insets outline the rightward loop of the embryonic heart tube in the control (A1), a reversed loop (B1) or no clear loop direction (C1) in the mutant samples. (A2-C2) Brightfield images of the head of the same individuals at E18.5. The red arrows point to cleft upper lips and the black asterisks to undetectable eyes in mutant fetuses. (A3-C3) Brightfield images of the hindlimb at E18.5, with numbered digits, showing polydactyly in the mutants. (A4-C7) Coronal (A4-C5) and transverse (A6-C7) sections of micro-CT scans, showing the lung and stomach (St) *situs* (A4-C4), the position of the heart apex in the thoracic cavity (red arrow in A5-C5), the position of the pulmonary venous return (red arrow in A6-C6) and the position of the inferior *vena cava* (blue arrow in A7-C7). (A8-C9), HREM images of the heart at E18.5, isolated from the same individuals. In coronal sections of the four cardiac chambers (A8-C8), the white asterisk shows complete atrioventricular septal defect in the mutants. In transverse sections (A9-C9), the relative positions of the aorta and pulmonary artery appear normal in the mutants. The white dotted line represents the plane of bilateral symmetry, bisecting the neural tube. Left and right anatomical features are annotated in blue and red respectively. Ao, aorta; Caud., caudal; Cran., cranial; L, left; LA, left atrium; Li, liver ; LLuL, left lung lobe; LV, left ventricle; PA, pulmonary artery; PCLuL, post-caval lung lobe; Pe, pericardial effusion; R, right; RA, right atrium; RMLuL, right middle lung lobe; RSLuL, right superior lung lobe; RV, right ventricle; St: stomach.

The external examination of E18.5 fetuses showed craniofacial malformations (Fig. 6A2-C2) and polydactyly (Fig. 6A3-C3) in 2/2 alive mutants, consistent with previous observations (Vierkotten et al., 2007). By micro-CT, we assessed laterality defects of visceral organs and found left isomerism of the lungs (Fig. 6A4-C4) and bronchi in both mutants. However, the laterality of other visceral organs was not affected in these cases. The liver and spleen were slightly smaller compared to controls (Fig. 6A5-C5, Movie 5). Congenital heart defects were evaluated from micro-CT and HREM. Both mutants showed complete atrio-ventricular septal defects (Fig. 6A8-C8), which precludes the use of valves as a criterium to identify the left and right ventricles. One mutant fetus had a normal heart *situs* (Fig. 6A5-B5), and normal ventricular anatomy, but an abnormal connection of the pulmonary veins to the right atrium, close to the inferior *vena cava* (Fig. 6A6-B7). The inferior *vena cava* appeared interrupted at the level of the liver and compensated by an azygos return of the right superior *vena cava* (Movie 5). In addition, the pulmonary artery branches were almost undetectable, suggesting hypoplasia (Movie 5). Thus, with a normal position of the morphologically left/right atria (S) and ventricles (D), and of the great vessels (S) (Fig. 6A8-B9), this mutant heart is described as {S,D,S} in the segmental approach. The other mutant fetus showed mesocardia (Fig. 6A5, C5) with an abnormal right aortic arch (not shown), as well as left isomerism of the pectinate muscles (Fig. 6A9, C9). The inferior *vena cava*, abnormally running on the left of the abdominal cavity (I, *situs inversus*), appeared dilated and entered the left part of the common atrium (Fig. 6A7, C7). We observed an abnormal insertion of the pulmonary vein collector in the right part of the common atrium (Fig. 6A6, C6). Thus, with a normal position of the morphologically left/right ventricles (D) and of the great arteries (S), this mutant heart is described as {I,D,S} in the segmental approach (Van Praagh, 1992).

By combining isomerism of the airways, together with atrial and venous return anomalies, the phenotype of *Rpgrip1l* mutants at E18.5 is heterotaxy, according to the criteria of (Lin et al., 2014). Using this mutant line, in which an abnormal embryonic heart shape correlates with congenital heart defects and isomerism of the airways, we validate the multimodality imaging pipeline to phenotype laterality defects in the mouse.

## Discussion

Based on sequential 3D imaging, the multimodality imaging pipeline that we have developed provides an extensive analysis of left/right anomalies, as well as congenital heart defects, using the nomenclature of paediatric cardiologists in the mouse. We provide annotated reference 3D images of asymmetric organ structures, including the embryonic heart loop at E9.5, visceral organs and great vessels at E18.5. The pipeline combines *in utero* analysis of embryonic shapes, by non-invasive micro-ultrasound imaging, with determination of the *situs* of visceral organs around birth, in their endogenous environment, by micro-CT. More resolutive assessment of congenital heart defects is provided by HREM. This standardised pipeline, which was validated in a mouse model of heterotaxy, offers a cost-effective, fast and easy-to-implement framework for phenotyping mouse mutants at multiple scales and multiple stages.

### Limitations and future extension of imaging modalities

Our pipeline combines advantages of 3 imaging modalities. Imaging organogenesis in the mouse embryo in a non-invasive manner is a challenging issue. Here, we focused on the embryonic heart tube which has been thoroughly quantified in fixed dissected samples (Le Garrec et al., 2017). With high frequency micro-ultrasound imaging, we were now able to assess the overall shape of the heart loop of individual embryos through the pregnant mother, and correlate, in *Rpgrip1l* mutants, abnormal looping of the embryonic heart with congenital heart defects. It will be possible to extend the analysis to other organs and other stages, without any detected adverse effects on *in utero* development. Micro-ultrasound imaging is less resolutive than OCT (30-100 μm versus 2-10 μm) (Lopez et al., 2015; Syed et al., 2011), but it has a higher imaging depth and so can be used at earlier stages, when the deciduum is thicker (E9.5 versus E12.5). It is also much less invasive compared to the required externalisation of the uterine horn for OCT. With technical development of the probe performance, the resolution of micro-ultrasound imaging is expected to further improve. Emerging methods, such as photo-acoustic microscopy, in which acoustic detection can reach a high imaging depth with a resolution varying in depth between 10-150 μm, are promising alternatives for *in utero* imaging, but limited to the cardiovascular system, because of the stronger optical absorption of haemoglobin (Laufer et al., 2012).

To perform non-destructive imaging of internal organs in mouse mutants, micro-CT has been developed (Degenhardt et al., 2010; Dickinson et al., 2016; Wong et al., 2014), given its fast speed of imaging, high resolution, and using lugol as a relevant contrast agent to visualize soft tissues such as the heart and vasculature, and also liver, lung and intestines. In addition to qualitative description of anatomical features, this technique is relevant to morphometric quantifications. Micro-CT has the advantage to be faster and less expensive than MRI (Magnetic Resonance Imaging) (Degenhardt et al., 2010; Norris et al., 2013), and has a wider field of view compared to OPT (Hsu et al., 2016; Sharpe et al., 2002). A limitation of this technique comes from adverse effects of the contrast agent, which induces slight tissue swelling (Degenhardt et al., 2010) and artefact aggregates during pre-treatments for HREM. This limitation, potentially affecting image segmentation, did not interfere with the resolution of the asymmetric shape and position of internal organs.

HREM offers an unprecedented histological resolution with 3D rendering, to resolve fine anatomical left/right variations. This technique, which is destructive, is used as a final step in the pipeline. It is relevant to morphometric analyses, as well as quantification of trabeculation or myofibre orientation (Garcia-Canadilla et al., 2018; Paun et al., 2018). HREM was used here for the heart, but could be extended to other explanted organs, or the whole fetus (Geyer et al., 2017).

### Differences in organ asymmetry between the mouse and human

Our annotated 3D images and organ segmentations provide a comprehensive reference of asymmetric organs in the mouse. Left-right features in the mouse are mainly similar to that in the human, with some exceptions (Table 1). The bronchus anatomy is the same with an earlier subdivision of the right bronchus compared to the left, whereas the number of lung lobes is different: 4 on the right in the mouse, compared to 3 in the human, and 1 on the left compared to 2 in the human. In the right lung, the post caval lung lobe is specific to the mouse (Thiesse et al., 2010). Another lobulated organ in the mouse is the liver, which is not the case in the human (Fiebig et al., 2012). In the human, the liver is a single continuous structure, subdivided in two regions on either side of the falciform ligament, conveniently named left and right liver lobes. This is equivalent in the mouse to the medial liver, which is also continuous between the left (LMLiL) and right (RMLiL). However, the mouse liver has additional left (1) and right (3) lobes. In the mouse as in the human, the stomach and spleen are localised on the left side of the body. The colon forms a leftward loop in the human, whereas in the mouse, it has a short C-shape curvature, going dorsally and on the left (Henke et al., 2018; Nguyen et al., 2015).

**Table 1.** Similarities and differences between human and mouse organ situs.

| **Anatomical criteria** |  | **Human** | **Mouse** |
| --- | --- | --- | --- |
| Differences | **Pulmonary venous return** | 4 pulmonary veins connected by 4 distinct ostia to the left atrium | 4 pulmonary veins connected by one collector to the left atrium |
|  | **Systemic venous return** | Regression of the left superior caval vein. Independent and thin coronary sinus. | Persistent left superior caval vein, connected by a dilated coronary sinus to the right atrium |
|  | **Lungs** | Right lung: 3 lobes<br>Left lung: 2 lobes | Right lung: 4 lobes<br>Left lung: 1 lobe |
|  | **Liver** | 2 virtual left/right lobes | 2 virtual left/right medial lobes + 3 right lobes + 1 left lobe |
|  | **Colon** | Leftward loop | Dorsal loop, on the left side |
| Similarities | **Heart** | 4 cavities, with asymmetric left/right ventricles and asymmetric left/right atrial appendages, 2 arterial valves and 2 atrioventricular valves, aorta posterior and right to the pulmonary artery, left aortic arch, right-sided inferior *vena cava* |  |
|  | **Bronchi** | Right bronchus: early first division<br>Left bronchus: late first division |  |
|  | **Stomach** | Lateralised on the left side |  |
|  | **Spleen** | Lateralised on the left side |  |

The heart anatomy of the mouse is very close to that of the human. Differences are most obvious in the venous return (Webb et al., 1996). The left superior *vena cava* is persistent in the mouse, whereas it regresses in healthy humans. The coronary sinus lies in continuity with the left superior *vena cava* in the mouse, whereas it becomes blind in the human (Kaufman and Richardson, 2005). Thus, bilateral superior *vena cava* is a heterotaxy feature specific to humans. In the human each pulmonary vein is connected individually to the left atrium by an ostium, whereas in the mouse, there is a collector draining together the four pulmonary veins to the left atrium. The external shape of the atrial appendages is more asymmetric in the human, with a narrow finger-like left atrial appendage and a broad and blunt right atrial appendage (Jacobs et al., 2007). All other left-right features are similar in the human and mouse. The aorta, which is posterior and right compared to the pulmonary artery, is connected to the left ventricle, and the pulmonary artery to the right ventricle. These great vessels spiral around each other. The aortic arch is on the left side of the body, while the inferior *vena cava* is on the right side.

### Physiological, anatomical or embryological nomenclature of left/right features

When phenotyping laterality defects, an important question is the rationale used to name asymmetric organ segments as left or right. A physiological perspective predominates in clinical cardiology, to distinguish, in reference healthy patients, the right heart, driving the circulation of deoxygenated blood, from the left heart, driving the circulation of the oxygenated blood. In this perspective, the inferior and superior vena cava, right atrium and ventricle and pulmonary arteries are defined as right features (shown in blue in Figures 4–5), whereas the pulmonary veins, left atrium and ventricle and aorta as left features (shown in red in Figures 4–5). In some instances, this perspective differs from the anatomical location. The coronary sinus, which drains deoxygenated blood, is considered physiologically as a right structure, although it is connected to the left *vena cava* in the mouse. The aortic valve, which drains deoxygenated blood is a physiologically left structure, although it lies on the right of the pulmonary trunk.

In an embryological perspective, the rationale is focused on the origin of cardiac segments, from the pool of left/right precursor cells in the lateral plate mesoderm after gastrulation, which receive distinct molecular signatures, such as Nodal signaling or Pitx2 transcriptional modulation on the left side (Collignon et al., 1996; Furtado et al., 2011; Liu et al., 2002). The fate and lineage of left versus right heart precursor cells was traced with DiI labelling (Domínguez et al., 2012) and clonal analyses15,16. In the embryological perspective, the right superior caval vein, the right atrium and the aorta, which derive from right progenitors, are considered as right structures, whereas the left superior caval vein, left atrium, pulmonary veins and pulmonary trunk which derive from left progenitors, are considered as left structures (see Meilhac and Buckingham, 2018). Thus, for the aorta, pulmonary trunk and left superior caval vein, there are discrepancies between the physiological and embryological perspectives. This may also affect the coronary sinus, which lies in continuity with the left superior caval vein. The left atrium was shown to receive a contribution from both right and left precursors (Domínguez et al., 2012). The origin of the ventricles has not been demonstrated yet, although the process of fusion of the heart tube would suggest a double left/right origin for each ventricle, which is also supported by genetic tracing with the left determinant Pitx2 (Ai et al., 2006; Furtado et al., 2011; Liu et al., 2002). The embryological perspective thus differs from the anatomical location in the case of cardiac chambers. This is because the anatomical location of cardiac segments is the result of asymmetric morphogenesis, from the rightward looping of the heart tube (Le Garrec et al., 2017) or the rightward rotation of the outflow tract (Bajolle et al., 2006).

In summary, our multimodality imaging pipeline opens novel perspectives for an exhaustive phenotyping of mouse mutants and provides novel insight into the embryonic origin of laterality defects and the mechanisms of congenital heart malformations. Beyond cardiovascular research, applications of this novel pipeline can easily be extended to the study of asymmetric morphogenesis of other organs. The fast and standardised collection of images is relevant to automated and quantitative image analysis procedures, which are emerging.

## Methods

### Animal models

Control embryos were from a mixed genetic background. The *Rpgrip1l*^*+/−*^ mouse line (gift from S. Schneider-Maunoury) was maintained in a C57Bl6J genetic background. Embryonic day (E) 0.5 was defined as noon on the day of vaginal plug detection. Animal procedures were approved by the ethical committee of the Institut Pasteur and the French Ministry of Research.

### Micro-ultrasound imaging

Pregnant female mice were anaesthetized with 4% isoflurane (in oxygen) for induction and 2% for maintenance. The abdomen was shaved using a depilatory cream to minimize ultrasound attenuation. The animal was restrained on a heated platform with surgical tape, maintaining a normal body temperature during imaging. Heart rate, temperature and breathing were monitored with paw and rectal probes. Ultrasonic gel was applied on the skin to perform non-invasive transabdominal imaging, using the Ultra High-Frequency Imaging Platform Vevo2100 (Visualsonics) with a 50 MHz probe (MS-700). Fast scans of the uterine horns and of each embryo were acquired to identify the embryo position in the uterus. A 3D+t scan of each embryo (E9.5) was acquired across the deciduum. The dataset comprises 94 images with an axial and lateral resolution of 30 and 50 μm respectively. The motor has a step size of 32 μm.

### X-ray micro-computed tomography

Foetuses were recovered before birth, at E18.5, carefully monitoring their position in the uterine horns. They were euthanized by decapitation. The thoracic skin was removed to allow penetration of the contrast agent; the left arm was removed, as a landmark of the left side. Blood was washed out in phosphate-buffered saline (PBS) and the heart was arrested in diastole with 250 mM cold KCl. Samples were stained in 100% Lugol (Sigma-Aldrich) over 72 hours. Images of the thorax and abdomen were acquired on a Micro-Computed Tomography Quantum FX (Perkin Elmer), within a field of exposure of 10 mm diameter. The dataset, comprising 512 images of 20 x 20 x 20 μm xyz resolution for each sample, was analysed using OsiriX Lite software.

### High resolution episcopic microscopy

A control E9.5 embryo was dissected, incubated in cold 250 mM KCl, fixed in 4% paraformaldehyde and washed in PBS. E18.5 hearts were isolated after micro-CT imaging, washed in PBS to remove Lugol staining as much as possible, post-fixed in 4% paraformaldehyde and washed in PBS. Samples were dehydrated in series of methanol concentrations and embedded in methacrylate resin (JB4, Polysciences) containing eosin and acridine orange as contrast agents (Weninger et al., 2006). Single-channel images of the surface of the resin block were acquired using the optical high-resolution episcopic microscope (Indigo Scientific) and a GFP filter, repeatedly after removal of 1.56 μm (at E9.5) or 2.34 μm (at E18.5) thick sections. The dataset comprises 1,200 images of 1.45-2.43 μm resolution in x and y at E9.5 and 1,600 images of 2.3-3.8 μm resolution in x and y at E18.5.

### Image analysis

Image sequences acquired by micro-ultrasound imaging were post-treated to generate a cubic resolution and volume rendering using the Volume viewer plugin from Fiji (ImageJ). The volume was re-sectioned to generate standardised coronal views, independently of the orientation of the acquisition. 3D images acquired by micro-CT were segmented with the AMIRA software (Thermo Fisher Scientific). 3D images acquired by HREM were analysed using the Imaris software (Bitplane): the heart at E9.5 was segmented as described in11, surface rendering of the E18.5 heart and optical sections were obtained with the oblique slicer. Volume rendering of E18.5 heart was obtained with the Volume viewer plugin from Fiji (ImageJ), after adjustment of the resolution as cubic.

## Acknowledgments

We thank M. Weiss and S. Bernheim for helpful discussions, V. Benhamo for technical assistance, J-p.Mulon for assistance with movie formatting, I. Anselme and S. Schneider-Maunoury for providing *Rpgrip1l* mutants, the histology platform of the SFR Necker, S. Corroyer and the Ultrastructural BioImaging platform of the Institut Pasteur, J. Sadoine, L. Slimani, G. Renault and the Plateforme Imagerie du vivant of the Université Paris-Descartes for X-Ray micro-CT and ultrasound imaging.

